## Supplementary figures and images for "Cas9-Mediated Knockout of Ndrg2 Enhances the Regenerative Potential of Dendritic Cells for Wound Healing"

### Extended Data Fig 1

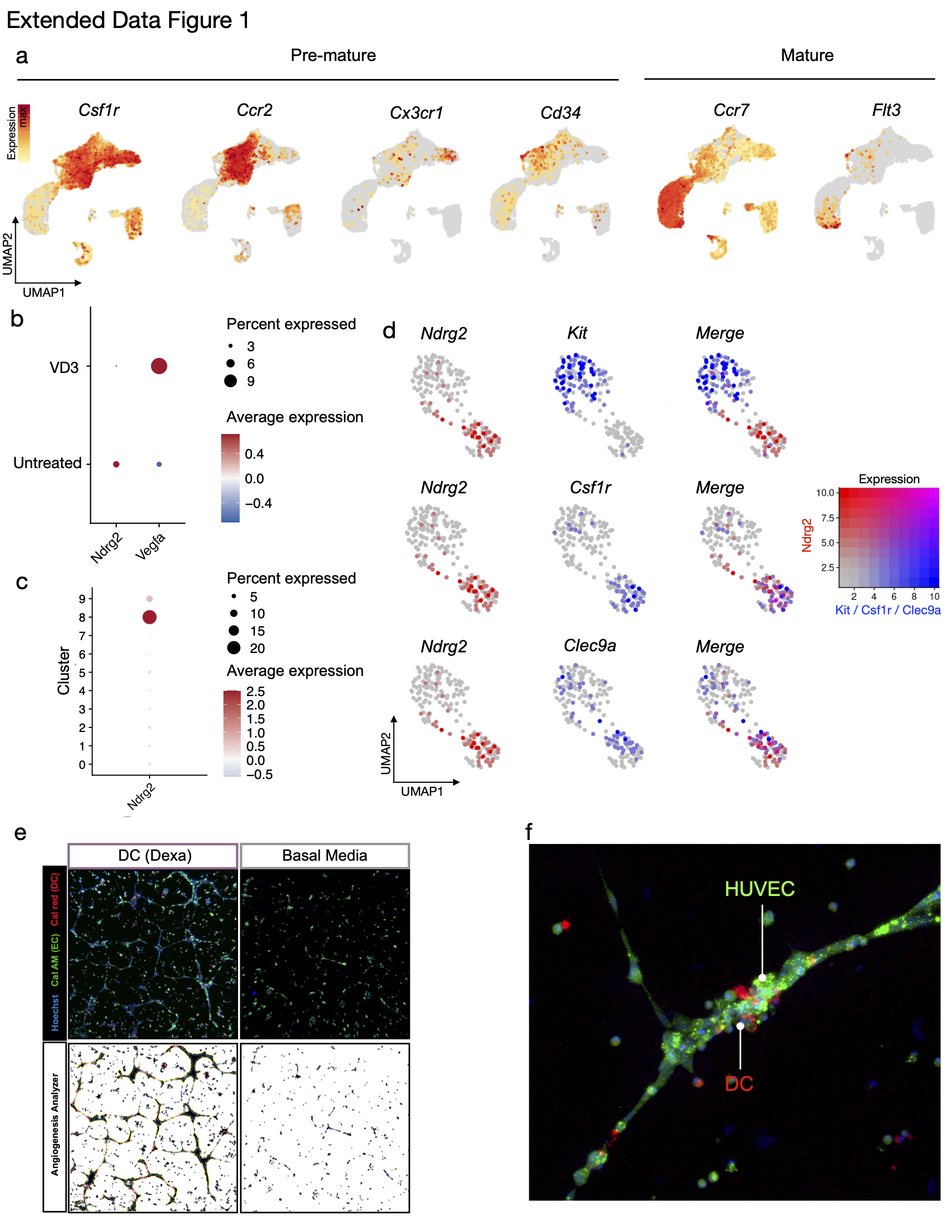

### Extended Data Fig 3

Extended Data Figure 3

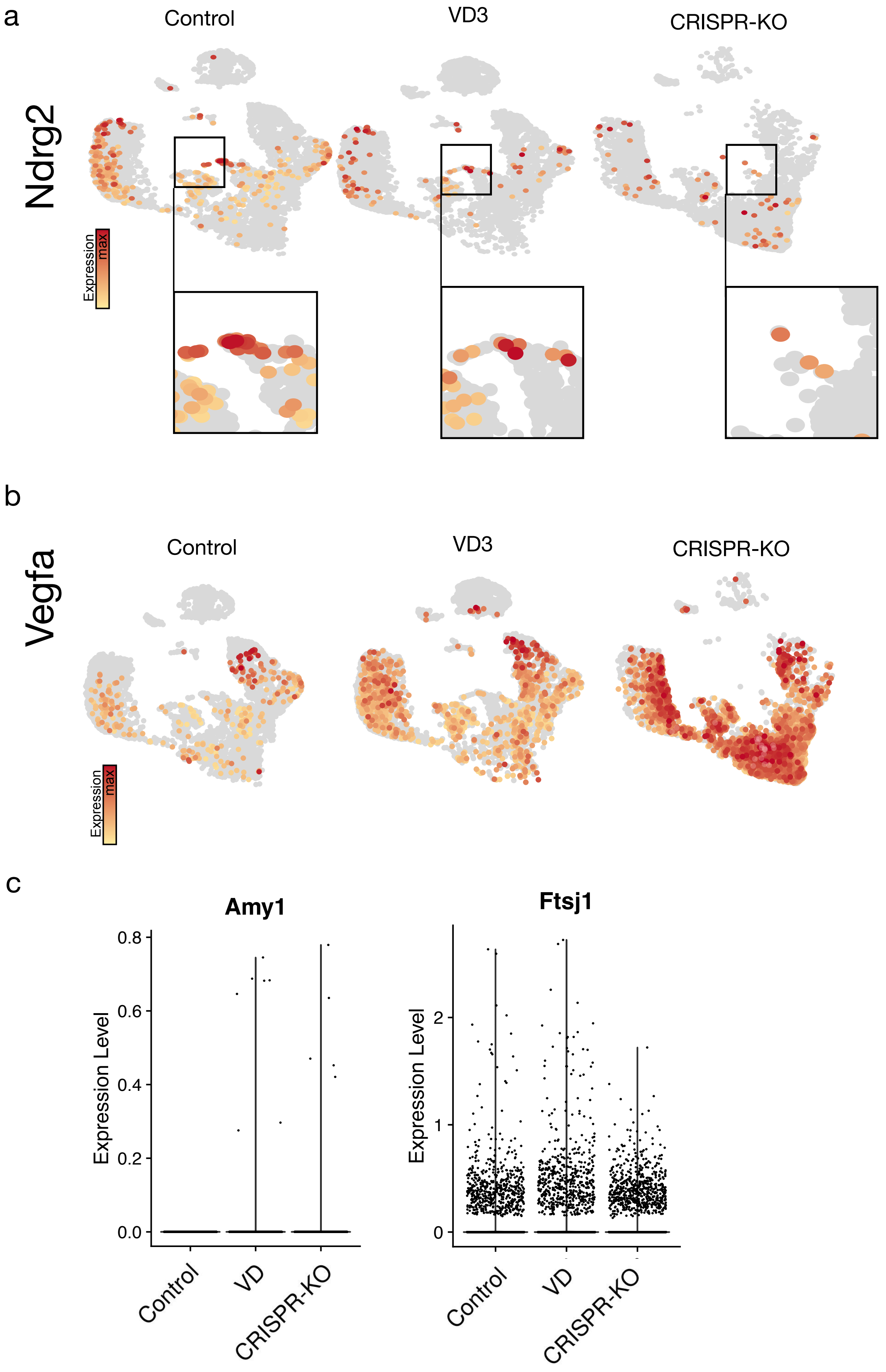

### Extended Data Fig 4

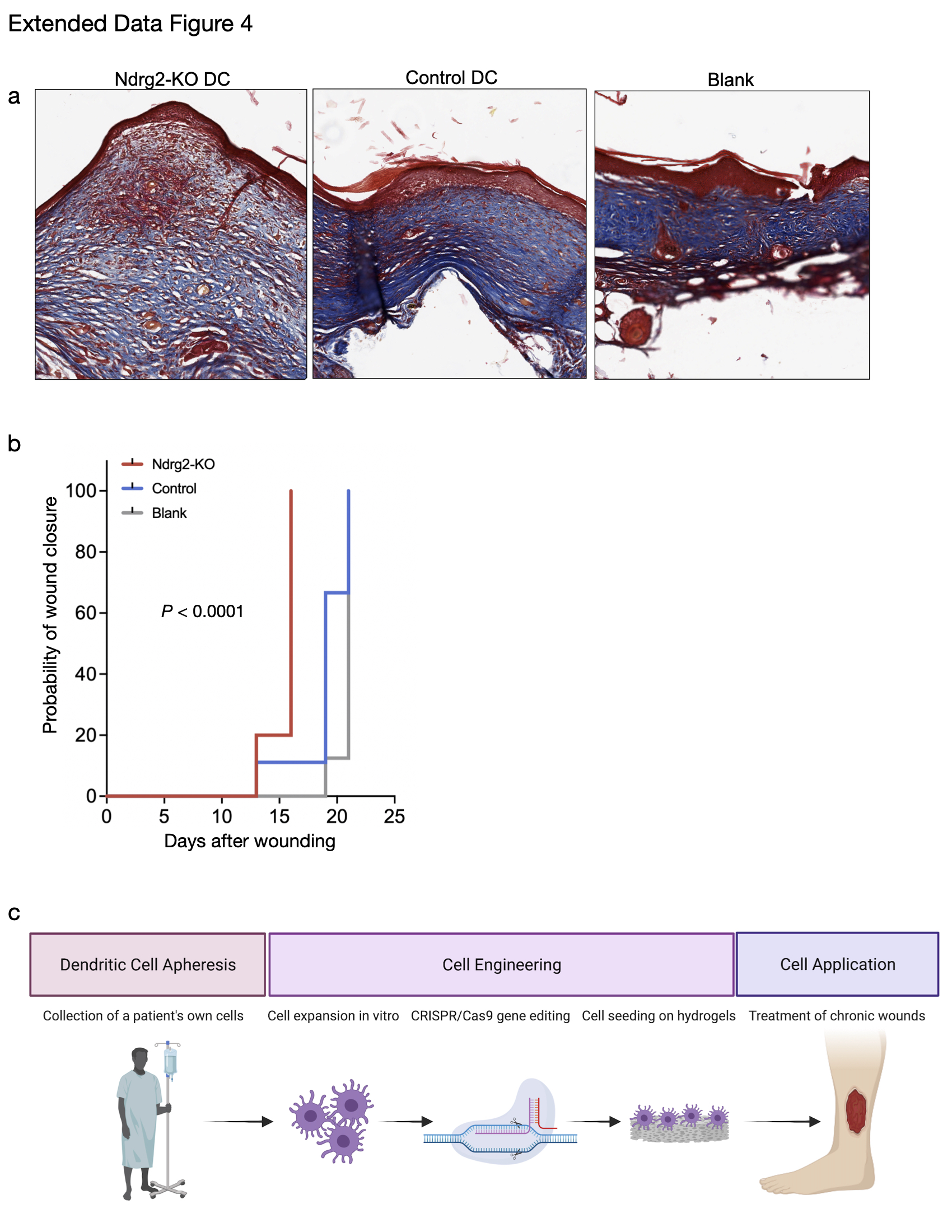
