## Extended Data Fig 2 for "Cas9-Mediated Knockout of Ndrg2 Enhances the Regenerative Potential of Dendritic Cells for Wound Healing"

Extended Data Figure 2

a

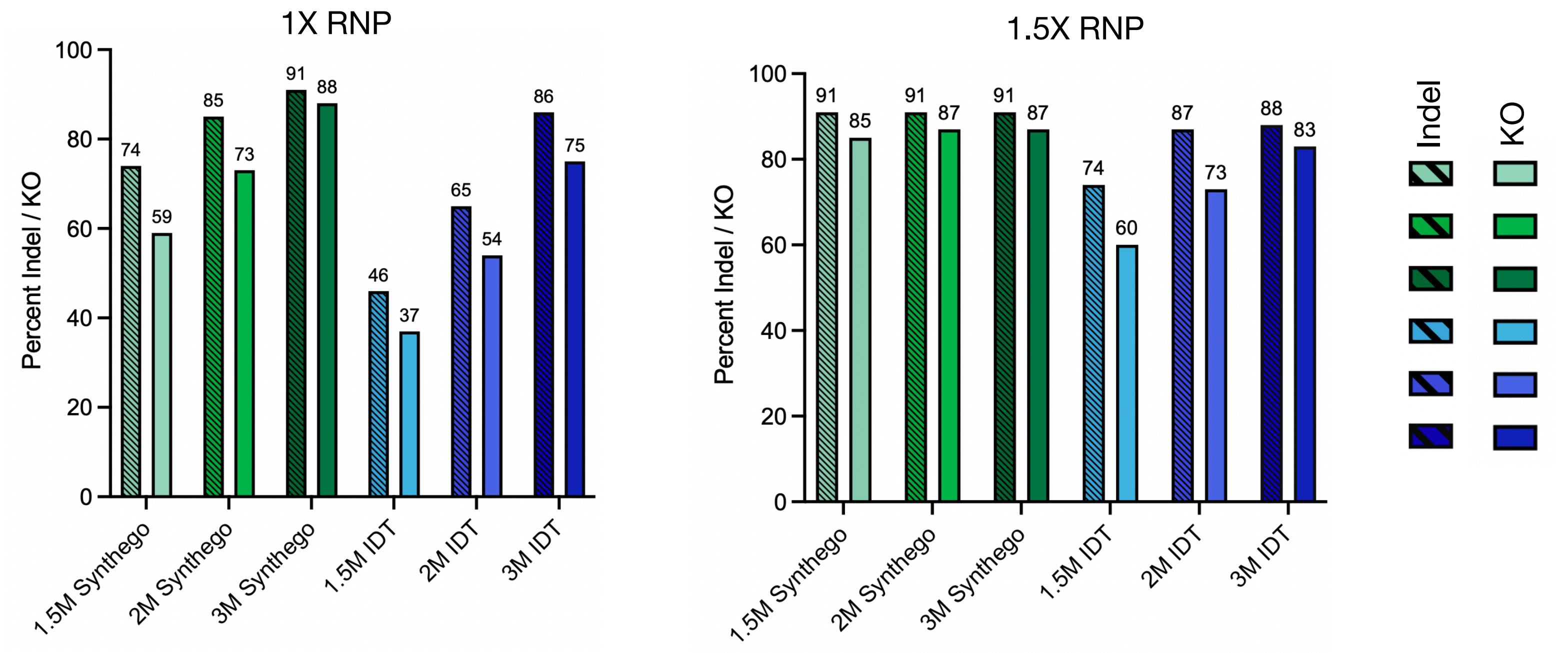


b Alignment between predicted off-target sites and sgRNA sequences

```
            chrX:126131778-126131801  -------------CCAGAAGAAACAAGTGAGATGCT------ 23
            chr15:32342483-32342507   -------ATGTGTAGTAGTATGGACCTGGG----------- 24
            chrX:145775172-145775196  -------GTGTTTAGTGCAATGTGACTGGGG---------- 24
sgNdrg2_2   Ndrg2+51911486            --------------------GCAGTCTCGGGTGTTTGTCC- 20
            chrX:116833564-116833587  --TTCCTTGATGTGCCAAGGTGGGG---------------- 23
            chr4:31726284-31726307    -CCCCCATGGCATAAGCACATGAG----------------- 23
            chr16:92812400-92812424   -----------------CCATGGGACCCATGTTCTTATCAT 24
            chr8:123766177-123766202  ------ATGTCCTCAGAGCTTAGGACCCAGG---------- 25
          → chr3:113572680-113572703  --------ATCTTCAGAGCTTGGCCCCGGAA---------- 23
sgNdrg2_1   Ndrg2+51911432            --------ATGTTCAGAGCATGGGACCG------------- 20
            chrX:8236822-8236846      CCGGACTGGCTAGAACTACAGGTG----------------- 24
            chrX:63795690-63795713    -CCCCTCCCCATGCCCTGAATACA----------------- 23
            chrX:60844817-60844840    CCACTGACATGCTCCCTACATGA------------------ 23
            chrX:48305848-48305871    -CCCCAGTCCCATCTCTGAAAGCC----------------- 23
            chrX:73054709-73054732    CCTCCTTCCCTCTCTCTGAACTC------------------ 23
            chrX:74953083-74953106    --CCTCTCTCTCTCTCTGAACATGG---------------- 23
            chrX:136367862-136367885  -----CCTCCATAGGA-TCGTGAGGATAA------------ 23
            chrX:52002777-52002802    CCTAGGTTCCATGACC-TCTGGAGAA--------------- 25
            chrX:11424965-11424988    ----CCCTGAAACACCCACAACTGTCT-------------- 23
          → chrX:96613063-96613088    -CTACATTGGAGTTCATTCATGGAGG--------------- 25
          → chr9:89597186-89597209    ---CTACTGAAAGCCTGCCCTGGAGG--------------- 23
sgNdrg2_3   Ndrg2+51911539            ---CTCCTGAAGTTCTGCCATGG------------------ 20
```

c Off-target varian classes

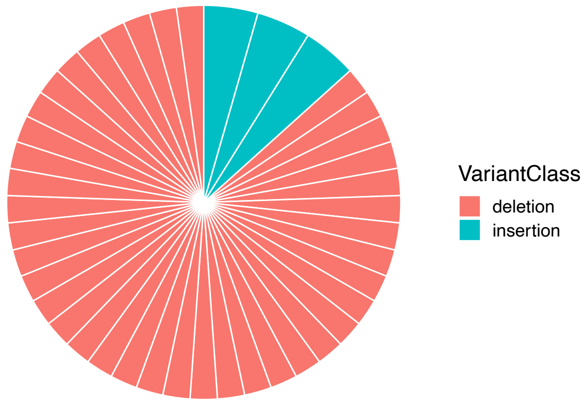
